# DNA Replication Stress Generates Distinctive Landscapes of DNA Copy Number Alterations and Chromosome Scale Losses

**DOI:** 10.1101/743658

**Authors:** Alice Mazzagatti, Nadeem Shaikh, Bjorn Bakker, Diana Carolina Johanna Spierings, René Wardenaar, Eleni Maniati, Jun Wang, Michael A. Boemo, Floris Foijer, Sarah Elizabeth McClelland

## Abstract

**Background:** We previously showed that a major driver of cancer chromosomal instability (CIN) is replication stress, the slowing or stalling of DNA replication. However, the precise drivers of replication stress in cancer and the mechanisms by which these cause CIN and influence tumour evolution remain unclear. Common fragile sites are well-known genomic locations of breakage after aphidicolin-induced replication stress, but their precise causes of fragility are debated, and additional genomic consequences of replication stress are not fully explored.

**Results:** Using single cell sequencing we detected DNA copy number alterations (CNAs) caused by one cell cycle under replication stress in diploid non-transformed cells. Aphidicolin-induced replication stress caused multiple types of CNAs associated with different genomic regions and features. Coupling cell type-specific analysis of CNAs to gene expression and single cell replication timing analyses allowed us to pinpoint the causative large genes of the most recurrent chromosome-scale CNAs. In RPE1 cells these were largely confined to three sites on chromosomes 1, 2 and 7 and generated acentric lagging chromatin and micronuclei containing these chromosomes. Different replicative stresses generated distinct profiles of CNAs providing the potential to interpret specific replication stress mechanisms from cancer cells.

**Conclusions:** Chromosomal instability driven by replication stress occurs via focal CNAs and chromosome arm-scale changes, with the latter confined to a very small subset of chromosome regions, potentially heavily skewing cancer genome evolution trajectories. Single cell CNA analysis thus reveals new insights into the impact of replication stress on the genome and provides a platform to further dissect molecular mechanisms involved in the replication stress response and to gain insights into how replication stress fuels chromosomal instability in cancer.

## Background

Most cancers exhibit chromosomal instability (CIN) that generates somatic copy number alterations, fuels tumour genome evolution, and is associated with poor prognosis. Replication stress drives chromosomal instability (CIN) in multiple cancer types^1,2^ and induced pluripotent stem cells^3^ and can be driven by oncogenes^4,5^, low nucleoside concentrations^6,7^ and difficult to replicate DNA sequences or structures^8,9^. High levels of replication stress can activate cell cycle checkpoints and lead to senescence that can form a barrier to tumour initiation^10,11^. Low levels however can bypass ATM and Chk1 activation^12^ allowing continued cell proliferation. Stalled forks induced by replication stress trigger multiple responses (reviewed in Ref 8^8^). It has also been shown that cells with under-replicated loci can trigger a measure of last resort mitotic DNA synthesis (MiDAS) to promote replication at these regions prior to completion of mitosis^13,14^. Failure to use any or all of these pathways may result in double strand breaks that are lethal to the cell if left unresolved. It has been observed that replication stress can induce lagging chromatin, anaphase bridges aneuploidy and micronuclei formation^15^. However, the processes that convert replication stress intermediates into large, chromosome-scale rearrangements, and whether other sub-microscopic genomic alterations also arise during replication stress remain incompletely characterised. The consequences of replication stress on the genome are only partially mapped, to date relying mainly on low-resolution cytogenetic analyses that detect breakpoint regions in metaphase spreads, termed common fragile sites (cFSs). Replication stress-induced DNA copy number alterations (CNAs) have also been detected using array comparative genomic hybridisation (aCGH)^16–19^ which provides high resolution but is limited to the detection of those CNAs that survive clonal outgrowth^16–18^, or appear frequently enough^19^ to allow their detection using bulk population analysis. Binding sites of the Fancomi Anemia protein FANCD2 following replication stress were also revealed using ChIP-Seq^20^. Lastly, MiDAS-Seq has mapped the genomic regions that frequently undergo mitotic DNA replication as a consequence of very late replication^21,22^. Replication stress induced CNAs identified so far may therefore represent only a minor fraction of the effects of replication stress on the genome.

To understand the precise and acute changes to the genome upon replication stress, we analysed CNAs induced in two diploid human cell types after one cell cycle under replication stress caused by low dose aphidicolin, using single cell low pass whole genome sequencing. This approach revealed multiple distinct classes of CNA, some of which were recurrent and clustered at characteristic genomic sites. A subset of large CNAs originated via chromosome breakage during mitosis. The resulting large chromosome fragments are mis-segregated at cell division, explaining the long-observed chromosome segregation errors caused by replication stress, and are incorporated into micronuclei, a well-established intermediate of chromothripsis^23^. Surprisingly, given the large number of previously mapped cFSs in the genome, this subset was confined to only a few sites in RPE1 cells, resulting in the majority of acentric chromatin fragments and micronuclei comprising material from only three chromosomes. Large CNAs were generally associated with large or giant genes, low surrounding gene transcription levels and late replication timing. One highly recurrent fragile region of chromosome 7 in RPE1 cells appeared dependent on RPE1-specific gene transcription of a nearby giant gene, AUTS2. A second method to induce replication stress (deletion of Mus81, a key endonuclease required for replication stress resolution) revealed a second, distinctive CNA signature characterised by focal losses. Altogether our study demonstrates that replication stress generates distinctive CNA spectra that reflect the specific origin of replication stressors. Determining the types and locations of genomic aberrations caused by deregulation of specific replication and repair factors also provides the platform to further study their precise cellular functions in maintaining genomic stability.

## Results

### Replication stress induced by low dose aphidicolin results in multiple classes of CNA that are associated with distinct genomic regions

To identify CNAs induced by replication stress we treated two diploid, telomerase-immortalised human cell types; retinal pigment epithelial (RPE1-hTERT, hereafter RPE1) cells and foreskin fibroblasts (BJ-hTERT, hereafter BJ), with low doses of the DNA polymerase inhibitor aphidicolin for 24 hours. Induction of replication stress and genomic instability was verified by increases in replication stress-related DNA damage, namely γH2AX foci in prometaphase cells, chromosome segregation errors in anaphase and 53BP1 bodies in G1 cells as well as micronuclei in interphase cells (Figure 1a-d; Figure S1a-l). We then performed low pass single cell sequencing of G1 cells to detect whole and partial chromosome copy number changes^24–26^ (Figure 1e) using EdU labelling to verify that cells in G1 had undergone DNA replication during that 24 h treatment period (Figure S2a,b). To gain greater depth of coverage we employed a modified library preparation protocol and sequenced at higher depth which generated ~ 0.04x coverage considering unique reads only (see Methods and Supplemental Table 1). This approach detected several classes of CNA, including a known sub-clonal trisomy of chromosome 12 and a clonal 10q amplification in RPE1 cells, and confirmed the XY karyotype of the BJ cells (Figure 1f-g; Figure S2c,d). In both cell types a set of recurrent focal amplifications were observed in both control and aphidicolin-treated cells. We reasoned that these likely represented existing structural variations present at clonal or sub-clonal frequencies since they were unique to each cell type and thus unlikely to represent a sequencing artefact. To verify this and benchmark the resolution of CNA detection using single cell sequencing we performed a SNP 6.0 array which verified the presence of a ~1.6 Mb focal amplification at chromosome 3p14.2 present in the majority of RPE1 cells (Figure S3a-d), in line with a previous single cell sequencing approach^27^. A higher resolution copy number analysis using 40 kb binning (see Methods) was able to detect smaller *bona fide* CNAs (~500 kb) that were also visible from SNP 6.0 array data, with higher accuracy of breakpoint position (Figure S3c,d). However a large number of likely artefactual CNAs were also observed with this analysis in both DMSO and aphidicolin-treated cells. Therefore we maintained the original (500 kb binning) resolution analysis to identify *bona fide* CNAs, but refined the position of these using the 40 kb binned analysis in order to pinpoint CNA breakpoints with the highest accuracy (Figure S3e; see Methods). To visualise the landscape of replication stress-induced CNAs we removed clonal copy number variants, and other CNAs that occurred in or near centromeric regions that were likely mapping artefacts (see Methods), and collated CNA events from 332 single RPE1 cells and 174 BJ cells (Figure 1h; Figure S2c). Aphidicolin treatment lead to an elevated rate of CNAs in both RPE1 and BJ cells with 38 % RPE1 cells and 44 % BJ cells exhibiting at least one aphidicolin-induced CNA (hereinafter ‘aCNA’) (Figure 1i). We noticed that aCNAs were either focal amplifications or deletions between 1-20 Mb, or much larger amplifications or deletions (>20 Mb) that extended from the single breakpoint to the end of the chromosome arm (‘large terminal aCNAs’; Figure 1j-l). Intriguingly, these distinct classes of aCNA tended to occur at different regions, and were differently distributed across the genome in both RPE1 and BJ cells (Figure 1g,h; Figure S2c,d), and we hypothesised that different mechanisms may operate to drive these classes of lesion.

**Figure 1:**
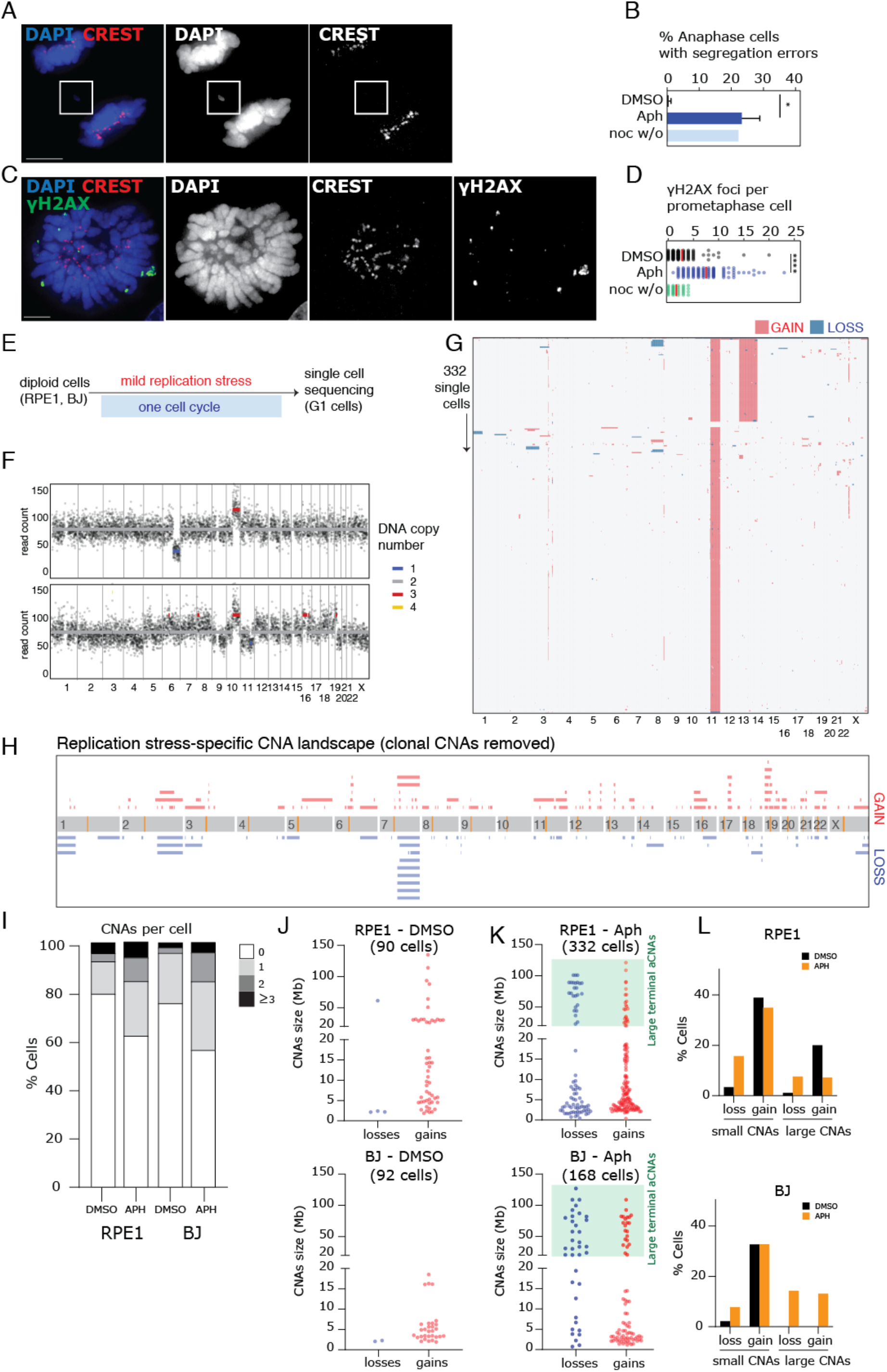
Replication stress induced by low dose aphidicolin results in multiple classes of CNA that are associated with distinct genomic regions. (**A**) Immunofluorescence image of RPE1 anaphase cell with lagging chromosome. CREST antibodies were used to stain for centromeric proteins. Scale bar in this and all subsequent microscopy images represents 5 μM. (**B**) Segregation error rates in RPE1 anaphase cells after indicated treatments (summary of three experiments for 24 h DMSO and 24 h aphidicolin, one experiment for 4 h nocodazole with 1 h release; n=167, 150 and 36 cells, respectively). Red bar indicates mean and statistical test was an unpaired *t*-test. (**C**) Image of RPE1 prometaphase cell; DNA damage foci detected using γH2AX antibody. (**D**) Quantification of DNA damage in RPE1 prometaphase cells after indicated treatments (n=110, 105 or 37 cells, respectively). Statistical test was an unpaired *t*-test. (**E**) Schematic explaining experimental setup to generate Replication Stress CNAs. (**F**) Example of Single cell sequencing readout. Coloured bars represents average copy number for those reads. (**G**) Single cell sequencing data from 332 RPE1 cells treated with aphidicolin; blue represents copy number losses, red represents gains. Amplification of 10q and recurrent aneuploidy of chromosome 12 are known aberrations in the RPE1 line. (**H**) Diagram summarising all RPE1 aCNAs taken from (G) after removing clonal events (see **Methods**). (**I**) Quantification of frequency of CNAs in DMSO and aphidicolin treated RPE1 and BJ cells (n= 90, 332, 92 and 168 cells, respectively). (**J, K**) Distribution of aCNAs divided by size and by gains versus losses in RPE1 and BJ cells in DMSO (J) and Aphidicolin (K). Box highlights large terminal CNAs (>20 Mb) which span the chromosome arm up to and including the telomere. (**L**) Frequency of each CNA class as indicated.

### Analysis of genomic features of CNA sites reveals associations with late replication timing and proximity to large genes

In order to define the aetiology of aCNAs we searched for associations with genomic features previously linked with cytogenetically identified fragility. Replication timing of the genome is an ordered process with most regions of the genome consistently being replicated at a specific time window during S-phase^28,29^. Fragile sites have been associated with both late (common fragile sites^16,30–32^) and early replicated (early-replicated fragile sites^33^). Replication timing programs can vary between cell types, and therefore we performed single cell sequencing-based replication timing analysis in a similar manner to recent studies^34,35^ on RPE1 and BJ cells in order to precisely analyse the relationship between CNA position and specific replication timing. We isolated cells from 4 separate S-phase fractions (Figure 2a) before single cell sequencing to determine copy number profiles (Figure 2b) providing, to our knowledge, the highest resolution replication timing profile of human cells using single cell sequencing to date. Replication timing profiles from single cells were similar to replication timing profiles derived from sequencing of bulk-populations (Figure S4a). This provided the opportunity to define the replication timing based on the proportion of cells that had replicated a given 1 Mb genomic region as a sum score across the four quartiles of S-phase (‘Replication timing factor’, see Methods). We divided CNAs into losses or gains, and either ‘small’ (<20 Mb) or ‘large terminal’ (>20 Mb and extending to the telomere). In RPE1 cells, large terminal losses displayed replication timing that was significantly later than a set of randomly placed 1 Mb CNAs that serve as an *in silico* control (see Methods; Figure 2c). In BJ cells a bimodal distribution was observed with half the large loss breakpoints falling in the latest replicated quartile. Small aCNA breakpoints in both RPE1 and BJ cells also followed an apparent bimodal distribution, occurring mainly in either early/early-mid, or late S-phase (Figure 2c) suggesting these aCNAs could be associated with both common (late) and early replicating fragile sites.

**Figure 2:**
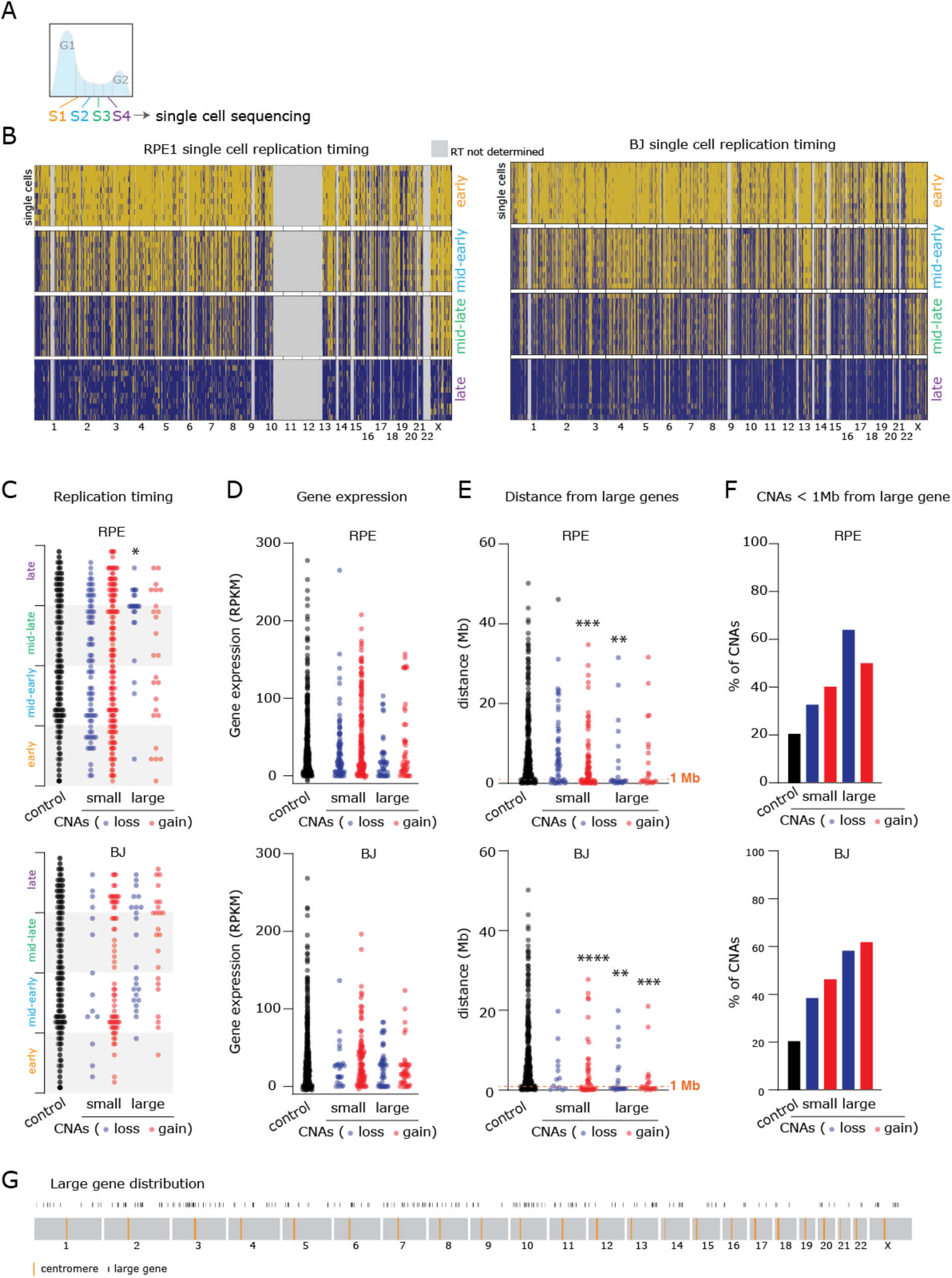
Genomic features of aCNAs. (**A**) Schematic indicating S-phase fractions isolated using FACS for single cell replication timing. (**B**) Single cell replication timing analyses for RPE1 cells (left panel) and BJ cells (right panel) from each S phase fraction as indicated. Dark blue indicates replicated genomic regions. (**C**) Replication timing factor was plotted for each aCNA in RPE1 and BJ cells. (**D**) Summed gene expression within a 2 Mb window around each aCNA breakpoint, separated into CNA classes as indicated for RPE1 and BJ cells. Control data comes from 470 in silico randomly generated 2 Mb windows, of which 376 were analysable for replication timing. (**E**) Distances from individual aCNAs to the nearest large or giant gene. Control represents 432 randomly generated genomic co-ordinates. (**F**) Proportion of CNAs that fall within 1 Mb of large/giant gene. (**G**) Schematic of large gene position along the genome. Statistical tests compare all aCNA classes to in silico control CNAs using a one way ANOVA Kruskal-Wallis test with post-hoc Dunn’s correction.

Previous studies have suggested a role for large genic transcription units in triggering hotspots of CNAs^16^. We therefore performed RNA sequencing to determine the abundance of gene transcripts genome-wide in RPE1 and BJ cells treated with DMSO or aphidicolin. Total expression levels were broadly similar between the two cell lines, and in line with previous studies^16,36^, aphidicolin treatment did not obviously impact gene expression globally, or of large or giant genes specifically (Figure S4b). We then analysed the summed gene expression within a 2 Mb window centred around breakpoints categorised into each CNA class, comparing to gene expression of random *in silico* placed breakpoint windows as a control (Figure 2d). No CNA categories exhibited gene expression that was significantly different to control regions in either RPE1 and BJ cells (Figure 2d).

Large (>600 Kb) or giant (>1 Mb) genes have been previously associated with fragility under replication stress^32,37,38^ although the exact causes remain debated. For small gains and large terminal losses in RPE and BJ, and additionally large terminal gains in BJ cells, the distance to the nearest large gene was significantly lower compared to a set of randomly-placed control regions (Figure 2e). Moreover all CNA classes had higher proportions of CNA breakpoints that were less than 1 Mb away from the nearest large/giant gene (Figure 2f,g). Taken together these data show breakpoints of large terminal aphidicolin-induced CNAs are usually late replicated and in proximity to large or giant genes, aligning them with previously identified common fragile sites. Interestingly, small CNAs caused by aphidicolin were also associated with proximity to large/giant genes.

### Transcription of giant genes can underlie cell type-dependent susceptibility to large aCNAs

We noticed that the region of chromosome 7q that was highly prone to large terminal aCNAs in RPE1 cells was far less frequently altered in BJ cells (Figure 3a). Recurrent breakpoints in chromosomes 1 and 2 also followed this trend although to a lesser extent. To determine the likely aetiology of the highly recurrent CNAs in these regions and to understand why they were not as susceptible in BJ cells, we visualised the genomic features of these regions obtained from analyses above (Figure 3b). As seen for large terminal CNAs in general (Figure 2e,f), we noted that these recurrent large aCNA-associated fragile regions were often obviously very close to large/giant genes (Figure 3b; Figure S5). Late or inefficient replication combined with proximity to large genes predisposes to fragility in many cFSs studied^39^ (Figure 2e-g). However replication timing at large genes closest to the most recurrent CNAs was similar between both cell types (Figure 3c), suggesting this was not a major factor in RPE1-specific fragility of these regions. AUTS2 and MAGI2 are giant genes close to the RPE1-specific aCNAs on chromosome 7q (Figure 3b). Referring to our RNA-Seq analysis (see above) we noted that AUTS2 was expressed in RPE1, and not BJ cells (Figure 3b (zoom); Figure 3d), indicating that AUTS2 was the causative gene for chromosome breakage at this site, and that gene expression is a requirement for the fragility of this large-gene associated site, in line with a previous study^16^. However, there was no clear involvement of gene expression status at other large or giant genes within RPE1-specific recurrent large aCNA sites, and gene expression of many candidate genes was undetectable in both cell types (Figure 3d) suggesting other factors may contribute to fragility at these loci. These data show we are able to pinpoint the gene responsible for precipitating the majority of large terminal CNAs in RPE1 cells. Moreover, in the case of FRA7B, a common fragile site, we show that fragility is due to the presence of two distinct regions; AUTS2 is expressed only in RPE1 cells and leads to a highly recurrent breakage in this cell line. MAGI2 is expressed in both RPE1 and BJ is a hotspot for CNAs in RPE1 (2 large terminal deletions and 2 focal gains) and BJ (3 large terminal CNAs and 2 focal gains) cells (Figure 3b,d). We then plotted the position of large terminal CNA breakpoints relative to large/giant gene positions relative to their promoter. This revealed that all large/giant-gene-proximal CNAs tended to originate within the central region of the genes, rather than the 5’ or 3’ ends specifically, in line with a previous study examining FANCD2 binding^20^ (Figure 3e,f).

**Figure 3:**
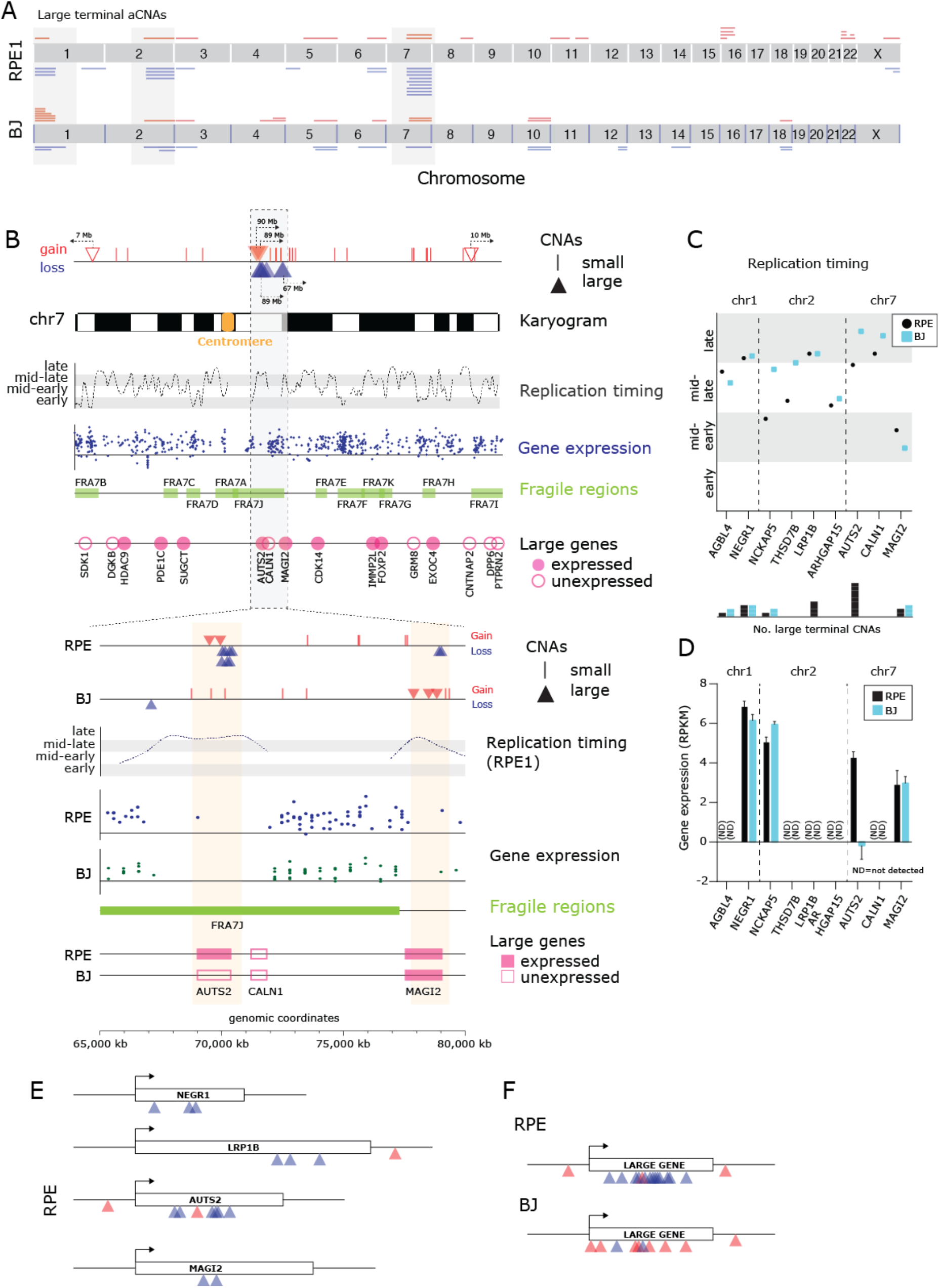
Transcription of giant genes can underlie cell type-dependent susceptibility to large aCNAs. (**A**) Schematic indicating locations and frequency of large, terminal aCNAs in RPE1 and BJ cells. (**B**) Schematic of chromosome 7’s genomic features. CNAs are marked (top) and aligned with a karyogram indicating the centromere and G-banding pattern, DNA replication timing from single cell CNA analysis, gene expression from RNASeq, fragile regions collated from the literature, and position of large (>0.6 Mb) and giant (>1 Mb) genes. Lower panel: Zoom of region containing 11 aphidicolin-induced large terminal chromosome losses and gains, indicating proximity of breakpoints to expressed giant genes AUTS2 and MAGI2, and their expression status in RPE1 and BJ cells. (**C**) Replication timing of large/giant genes nearest to recurrent large terminal aCNAs on chromosomes 1, 2 and 7 (**D**) Gene expression levels from both cell lines, of large genes nearest to recurrent large terminal aCNAs on chromosomes 1, 2 and 7. (**E**) Schematic indicating position of RPE1 large terminal aCNA events relative to most frequently affected large/giant genes and promoters. (**F**) Schematic indicating positional distribution of large terminal aCNAs in RPE1 and BJ cells to a normalised large/giant gene.

### A small number of genomic sites are prone to replication-stress induced chromosome mis-segregation events and micronuclei containing a small subset of chromosomes

We reasoned that aphidicolin-induced large terminal CNAs (Figure 3a) could be connected with chromosome segregation errors observed following aphidicolin treatment^1^ (Figure 1a,b; Figure S1h,i). Indeed, mis-segregating chromatin and micronuclei caused by aphidicolin lacked centromeres (‘acentric’) as judged by the lack of centromere-specific proteins or centromeric DNA sequences measured by immunofluorescence and fluorescence *In-Situ* hybridisation (FISH) respectively (Figure 4a,b; Figure S6a-e). Furthermore, the majority (78%) of lagging chromatin fragments contained a focus of DNA damage at one, or both ends, as marked using antibodies to γH2AX (Figure 4a,c). We reasoned that if chromosome mis-segregation in RPE1 cells resulted from the large terminal aCNAs initiated at these recurrent breakpoints, then these chromosomes should be overrepresented in lagging chromosome fragments. Because anaphases are rare in asynchronous RPE1 cells, we analysed the identity of chromosomal material encapsulated within micronuclei that result from mis-segregating chromatin thus providing a record of previous chromosome segregation errors. FISH-based painting of specific chromosomes revealed that 18, 12 and 42% of micronuclei in aphidicolin-treated RPE1 cells contained material from chromosomes 1, 2 or 7 respectively, whereas this enrichment was not observed in control cells (Figure 4d,e). This suggests that large terminal aCNAs are generated from breakage during mitosis, potentially as a result of condensation-induced rupture of under-replicated regions^40^, followed by either mis-segregation into the wrong daughter cell (leading to gain of genetic material), or incorporation into micronuclei (leading to loss of genetic material). Since micronuclei are lost during preparation for single cell sequencing^25^ this explains the bias towards losses of these large regions in single cell sequencing data (Figure 3a). These data are in agreement with a previous study that noted increased interphase aneuploidy of chromosomes harbouring common fragile sites in HCT116 and MRC5 cells^41^. Taken together with our findings above, replication stress caused by aphidicolin induces sub-microscopic, and microscopically visible chromosome evolution. Chromosome arm-scale CNAs originate at a very small subset of large/giant genes some of which must be expressed to induce fragility, leading to chromosome segregation errors and micronuclei.

**Figure 4:**
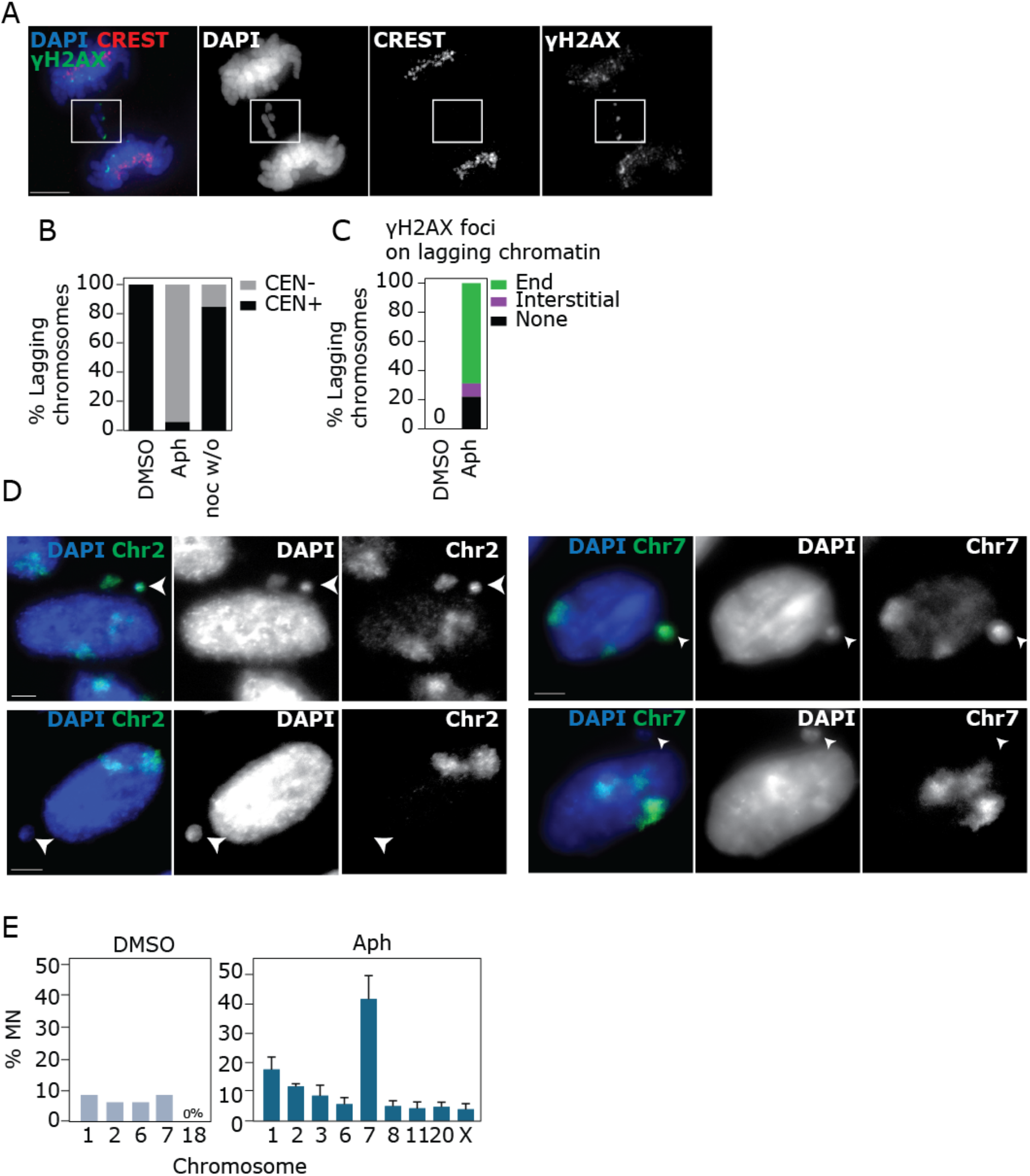
A small number of genomic sites are prone to replication-stress induced chromosome mis-segregation events and micronuclei containing a small subset of chromosomes. (**A**) RPE1 anaphase cell with acentric lagging chromosome; centromeric proteins stained with CREST, DNA damage foci detected by γH2Ax staining. (**B**) Centromeric status of lagging chromosomes in RPE1 cells after indicated treatments (24 h DMSO, 24 h aphidicolin, 4 h nocodazole 1 h release; n=2, 46 or 13 chromosomes, respectively). (**C**) Percentage of lagging chromosomes in RPE1 anaphase cells with DNA damage detectable on chromosome ends or within chromosome mass (n=35 chromosomes scored in aphidicolin). (**D**) Representative images of RPE1 cells stained with chromosome FISH paint to identify chromosomal identity of MN. (**E**) Quantification for positive staining of chromosome paints in RPE1 cells (DMSO one experiment, aphidicolin three experiments, 50-100 MN scored per chromosome per experiment).

### Depletion of Mus81 generates a distinctive profile of CNAs characterised by focal gains and losses

Next, we wondered whether distinct drivers of replication stress could be differentiated using single cell CNA analysis. Additionally we reasoned that knowledge of the position and genomic features of CNAs caused by specific replication stress mechanisms could give insights into their aetiology, in turn providing new information about the mechanism of action of specific components of the replication stress response. Therefore, we used siRNA to deplete Mus81, an endonuclease responsible for cleaving replication intermediates, and promoting mitotic DNA replication (MiDas)^13,14^, from cells using siRNA (Figure 5a). Depletion of Mus81 alone did not cause an increase in DNA damage, segregation errors, ultrafine anaphase bridges (Figure 5b-d) or micronuclei (Figure S7a-d). siMus81 treatment alone thus did not appear to cause microscopically visible loss of genome integrity. However Mus81 depletion in combination with aphidicolin treatment, led to increased DNA damage and UFBs compared to aphidicolin alone, in accordance with Mus81’s role in protecting against aphidicolin-induced genome instability^42^ (Figure 5d). Since aphidicolin treatment, in addition to large scale chromosome changes that were visible as chromosome segregation errors and MN, also induced a large number of other CNAs that would not be observable microscopically we wanted to assess whether Mus81 loss alone was capable of inducing cytogenetically invisible CNAs. We therefore sequenced single cells from siControl, and siMus81-treated cells (Figure 5e; Figure S7e). siMus81 treatment induced more CNAs per cell (Figure 5f,g), and these were enriched for focal gains and losses compared to siControl (Figure 5h). Moreover, siMus81-treated cells displayed a distinctive spectrum of CNAs when compared to aphidicolin treatment, comprising more interstitial focal losses and very few large terminal CNAs (Figure 5i, Figure S7d). We next analysed the genomic features of Mus81-induced CNAs as we did above for aCNAs. Here we classed small CNAs as less than 10 Mb given the spread of the CNA size. Replication timing and gene expression were not significantly different to *in silico* control regions (Figure 5j,k). There was an association with proximity to large genes that was highly significant for gains, both small (<10 Mb) and large (> 10 Mb) CNAs. However there was no significant difference in proximity to large genes between siControl and siMus81 CNAs (Figure S7f). Overall, these data reveal that Mus81 plays an important role in maintaining genomic stability at a subset of genomic regions, but that these regions do not appear to be enriched any of the classic fragility features, including late replication timing.

**Figure 5.**
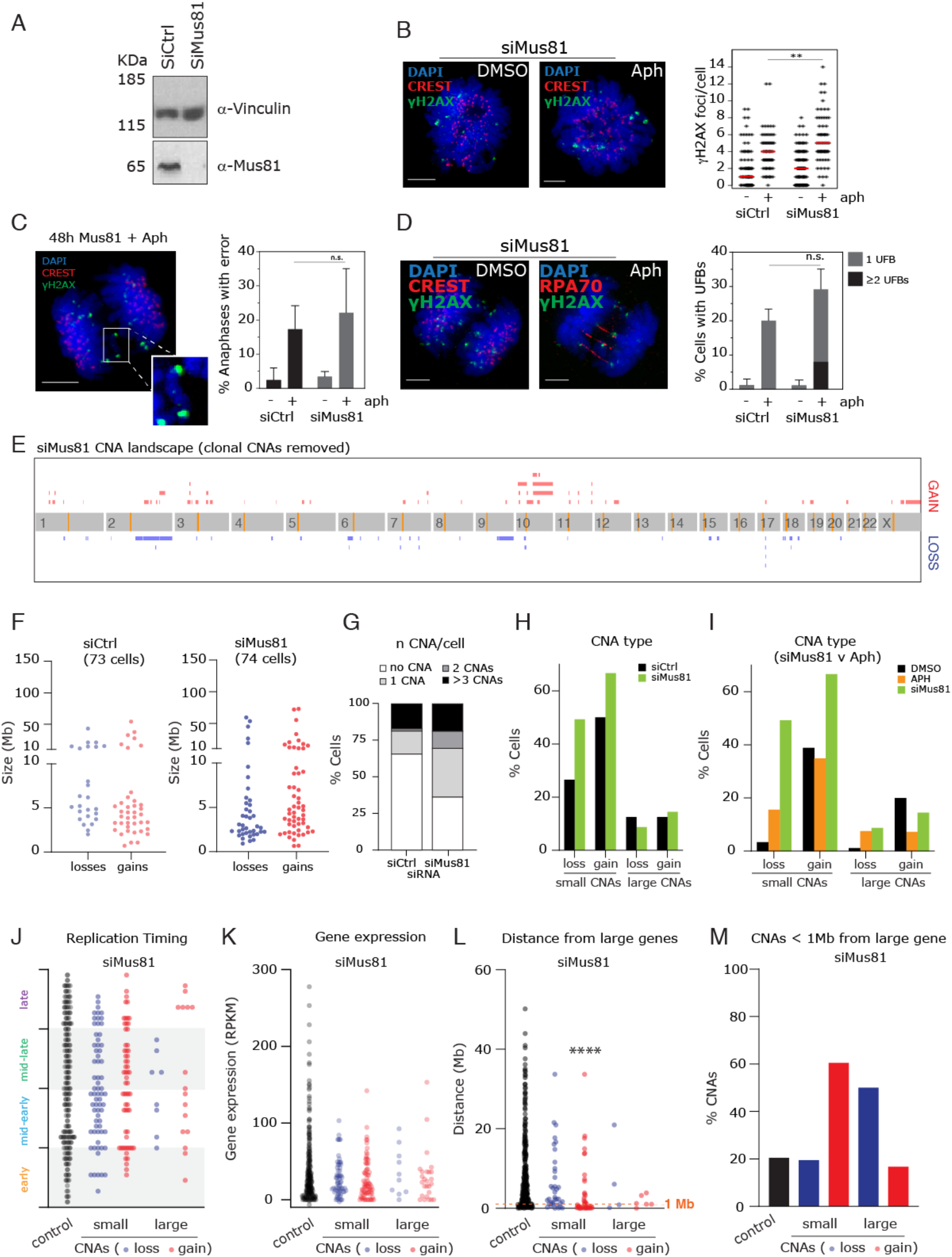
Depletion of Mus81 generates a distinctive profile of CNAs characterised by focal gains and losses. (**A**) Western blot to indicate loss of Mus81 protein in RPE1 cells after siRNA for 48 h. Vinculin was used as a loading control. (**B**) Immunofluorescence images of RPE1 cells stained with γH2AX to detect DNA damage in prometaphase cells treated with siControl or siMus81, and DMSO or aphidicolin as indicated. Right: quantification of γH2AX foci per cell. (**C**) Example image and quantification of chromosome segregation errors in anaphase in cells treated as indicated. (**D**) Example image and quantification of ultrafine anaphase bridges, marked using replication protein A (RPA70; red). (**E**) Diagram summarising all siMus81 CNAs taken from single cell sequencing of 74 cells (see heatmap in Figure S7) after removing clonal events (see **Methods**). (**F**) Distribution of CNAs divided by size and by gains versus losses following siControl (xx cells) and siMus81 (xx cells) treatment. (**G**) Frequency histogram of % cells exhibiting indicated CNA classes. (**H**) Frequency of each CNA class as indicated (**I**) Comparison between aphidicolin and siMus81 treatments for CNA type frequencies. (**J**) Replication timing factor was plotted for each CNA type as indicated. (**K**) Summed gene expression within a 2 Mb window around each siMus81 CNA breakpoint, separated into CNA classes as indicated. (**L**) Distance from large or giant genes was plotted for Mus81 CNAs. (**M**) Proportion of CNAs that fall within 1 Mb of large/giant gene. Control data in (J-L) from 432 randomly generated 2 Mb windows, as above. Immunofluorescence images are 25 or 30 z-stack projections. Scale bars 5 μm. Statistical tests in C and D are one-way ANOVA with post-hoc Dunnett’s test. Statistical tests in (J-L) compare all CNA classes to control using a one way ANOVA Kruskal-Wallis test with post-hoc Dunn’s correction.

## Discussion

### Single cell sequencing of replication stress induced CNAs provides a complement to previous studies and can detect non-recurrent focal amplifications

Here we sought to comprehensively assess the impact that replication stress has on the genome, motivated by a desire to understand the mechanisms that convert replication stress to genome evolution during cancer. We performed single cell sequencing to detect CNAs caused after one cell cycle and in the absence of any selective pressure that might influence the detection of CNAs formed by replication stress. We also analysed DNA replication timing at the single cell level, and bulk RNA sequencing to analyse gene expression in the same cell types, providing a complete picture of the factors involved in precipitating CNAs during replication stress from two different cell types. Previous studies tended to focus on a small number of recurrent fragile sites to understand molecular mechanisms operating to drive fragility under replication stress, potentially underlying some of the apparent conflicting views on the causes of fragility. A major advantage of our single cell sequencing approach is that we can follow not only large chromosomal breaks and gaps, but also the resulting copy number alterations induced at specific loci following replication stress at a resolution of 0.5-1.6 Mb. Replication stress-induced CNAs have been previously mapped to high genomic resolution using clonal outgrowth of cells where it was concluded that replication stress-induced CNAs tend to occur at cFSs^16^. Our approach is additionally able to detect CNAs that might not be propagated in the population during clonal growth. We showed that large terminal aCNAs, often arising close to large or giant genes, convert into lagging chromatin and micronuclei. These are likely to correspond to conventional cFSs. Additionally aphidicolin induced small losses and amplifications, many of which are unlikely to correspond to cFSs. First, many were not in recurrent positions and therefore would not be detected as common fragile sites (usually defined as representing at least 1% of cFSs). Moreover, small amplifications tended to occur in locations distinct from sites of large CNAs, suggesting they are caused, or resolved, by mechanisms distinct from those leading to previously-mapped cFSs.

### What is special about recurrent sites of large terminal CNAs?

Large and giant genes have been proposed to cause fragility under replication stress due to a number of mechanisms including late^16,30–32^ or delayed replication timing^36,39^, lack of origin density^43^, replication-transcription collisions during S-phase^5,16,44^ and gene expression in G1 that removes licensing complexes and reduces origin density^39,45,46^. We showed that large terminal aCNAs were generally close to large or giant genes, and that in the case of the most highly recurrent site of aCNAs in RPE1 cells this was likely due to the RPE1-specific gene expression of the giant gene AUTS2. Although the association between large genes and fragile regions is well established, pinpointing the exact gene(s) and mechanisms responsible for fragility in each fragile site (often spanning very large regions), has not been possible with cytogenetic techniques. Our single cell sequencing approach was able to identify two separate giant genes within (or just adjacent to) the common fragile site FRA7J that were recurrent sites of large terminal aCNAs. By comparing the position of breakpoints relative to position along large/giant genes we showed that breakage occurs within the gene bodies, although higher resolution sequencing analyses will be required to finely pinpoint these positions. Late replication timing or steady state gene expression did not readily explain all cell type-dependent differences in fragility at recurrent large aCNAs (Figure 3b-d) however. We also note that there are numerous expressed large genes residing within previously identified common fragile sites that do not precipitate CNAs following aphidicolin treatment (for example see chromosome 7 in Figure 3b and chromosomes 1 and 2 in Figure S5), suggesting other factors may contribute to the extreme bias towards fragility of the recurrent large terminal CNAs observed in this study. Furthermore causative factors could vary between specific fragile sites, highlighting the importance of assessing causative features across numerous fragile sites in a genome-wide manner.

We were surprised by how few sites were prone to large terminal aCNAs, despite the presence of approximately 150 large or giant genes in the genome, many of which are expressed, and tend to replicate late (Figure 3b)^32^. There may be other factor(s) involved in predisposing these specific genomic sites to fragility. For example these genes may lie in regions that are replicated early or mid S-phase in normal cells but could be delayed in timing in the presence of aphidicolin^36,39,47^ which could explain their fragility. In our study we analysed replication timing from DMSO-treated cells but it would be interesting to determine whether fragility of these regions correlates with aphidicolin-specific delays in replication timing. Our ability to characterise single cell-level differences would also provide interesting insights into the variability of such delays across individual cells. The timing or rate of active gene transcription may also be an important factor that we were unable to detect with RNA sequencing. Performing alternative methods to measure active gene transcription is likely to shed further light on the relationships between gene expression and CNA positions. A second possibility is that variation in the population at the single cell level could account for fragility, since although these regions are considered recurrent they still only occur in 1-5% cells. Gene expression is known to vary widely between individual cells. Therefore single cell RNA sequencing may give an indication of whether, despite similarities at the bulk population level, variability in gene expression at the single cell level could explain why a subset of single cells are prone to specific aCNAs.

### How does aphidicolin cause focal CNAs?

We did not identify obvious features of the surrounding genomic regions that would readily explain why small losses and gains would be induced by aphidicolin treatment, apart from a proximity to large genes. Small CNAs may occur not as increased predisposition to fragility *per se*, but an increased propensity to be aberrantly repaired or dealt with under replication stress. For example underlying sequence features such as tandem duplications may promote aberrant recombinatorial repair. In terms of small deletions, these might be intuitively suspected to be caused simply by under-replication. However these regions did not associate exclusively with very late replication timing as would be predicted from this. It is therefore tempting to speculate that these focal aCNAs may represent a novel class of lesion induced by replication stress that has hitherto gone undetected. Determining the molecular mechanisms rendering these positions fragile, and the DNA repair and bypass mechanisms involved in their generation is therefore likely to provide new insights into the processes connecting replication stress to chromosomal instability. A key step in completing the puzzle of the aetiology of replication stress-derived CNAs will be resolving the copy number breakpoints and any associated structural genomic variations at base-pair resolution, which will provide important clues to the molecular pathways connecting replication stress to genomic instability. In turn such analyses will be important to discover the aetiology of disease-associated copy number and structural variations currently being uncovered at rapid rates from genome sequencing studies^48,49^.

### Different replication stressors impact the genome in distinctive ways that can inform mechanisms of replication stress

Having identified a distinctive and informative set of CNAs following aphidicolin treatment, we were motivated to discover whether different insults to proper completion of DNA replication could also generate distinctive CNA spectra. In this study we depleted Mus81 and analysed CNAs in the absence of exogenous replication stress. Despite no obvious cellular responses in terms of DNA damage or segregation errors, removal of Mus81 from cells produced a distinctive CNA landscape, many of which were focal CNAs. These tended to occur very close to large or giant genes, but did not exhibit obvious differences in replication timing or overall gene expression. Interestingly, full depletion of Mus81 from cells, although leading to an elevated rate of aphidicolin-induced UFBs per cell, did not reduce the rate of acentric anaphase segregation errors or micronuclei caused by aphidicolin, suggesting that in RPE1 cells Mus81 is not essential for conversion of fragile sites into chromosome breaks as was previously shown in U2OS (osteosarcoma) and GM00637 (fibroblast) cells^50^. This could also be reflective of different fragile sites becoming expressed due to different mechanisms such as rupture during chromosome condensation upon mitotic entry40. Therefore the precise pathway leading to rupture and breakage at the three most common sites of large terminal deletion in RPE1 cells is not yet clear. In addition to our study, additional mechanism-specific CNA profiles have been revealed from murine models of cancer. Array CGH and single cell sequencing of tumours formed in mouse models of deregulated Mre11 (DNA double strand break repair factor) and the DNA replication licensing factor MCM2 revealed distinctive CNA landscapes biased towards small deletions, although these were necessarily the product of both mutation and selection during tumour evolution^51,52^. Single cell sequencing of p53 null RPE1 cells following Cyclin E1 or CDC25 also revealed distinctive CNA landscapes, characterised by focal CNAs and large terminal CNAs respectively^53^. Discovering the precise causes of CNAs caused by aphidicolin and Mus81 loss, in addition to those caused by other replication stressors, will likely uncover new insights into the mechanisms of specific components of the replication stress response.

### Replication stress could act to drive non-random CIN early in disease

Our data shows replication stress generates small CNAs that we identified from G1 cells following faulty S-phase, and thus appear not to activate cellular checkpoints, suggesting replication stress could act as a ‘stealth’ CIN mechanism in the presence of functional cellular checkpoints and challenging the view that overcoming DNA damage-induced cellular senescence is a necessary step for CIN and tumour formation^54^. Further analysis of the sizes and locations of CNAs that do, or do not elicit cell cycle arrest and senescence will be an important route to determining the potential for replication stress to act as an early driver of CIN and tumourigenesis without the need to overcome normal cellular checkpoints. Moreover the observation that in RPE1 cells locations of large, chromosome arm-scale events were essentially confined to three discrete loci affecting only chromosomes 1, 2 and 7 suggests that replication stress has the potential to heavily shape the evolution of tumour genomes in a non-random manner in a similar manner to our previous observations of non-random chromosome segregation caused by mitotic defects^25,55^.

## Ethics approval and consent to participate

Not applicable

## Consent for publication

Not applicable

## Availability of data and materials

All raw sequencing data will be accessible via the European nucleotide database upon publication

## Author contributions

NS and AM designed and performed the majority of cell biological experiments and data analysis supervised by SM. BB, RW and DCJS performed and analysed single cell sequencing data supervised by FF. EM analysed RNA sequencing data supervised by JW. MB wrote scripts for the analysis of clonal CNAs and visualisation of CNA landscapes. SM conceived the study, designed experiments, analysed data and wrote the manuscript with input from all authors.

## Acknowledgments

We would like to thank Susana Godinho for kind gifts of reagents. We thank Petter Larsson for generating python scripts to extract genomic features from publicly available databases and Daniel Muliaditan for creating in silico control coordinates.

## Methods

### Cell Culture and RNAi

All cell lines were maintained at 37°C with 5% CO_2_. hTERT-RPE-1 cells were cultured in DMEM Nutrient Mixture F12 Ham (Sigma); BJ cells in DMEM high glucose (Sigma). Media for both was supplemented with 10% FBS and 100 U Penicillin/Streptomycin. RPE1 and BJ cells were subjected to STR profiling to verify their identity using the cell line authentication service from Public Health England. Replication stress was induced using 0.2-0.4μM Aphidicolin (Sigma) for 24hr. RNAi was achieved by transfection of cells for 48 hr with 20 nM small interfering RNA (siControl [D-001210-02] and siMus81 [CAGCCCUGGUGGAUCGAUAUU], Dharmacon) using Lipofectamine RNAiMAX (Invitrogen) and Optimem (Gibco). Medium was replaced with fresh media 24 hr after addition of siRNA, and either DMSO or aphidicolin added 48 hr after addition of RNAi for a further 24 hr.

### Western blotting

Cell lysates were prepared using lysis buffer (20mM Tris-HCl, pH 7.5, 1% Triton X-100, 150mM NaCl, 5mM EDTA, 50mM NaF, 1mM PMSF, protease inhibitors (Roche)). Immunoblots were probed using antibodies against Mus81 (M1445, Sigma Aldrich) or Vinculin (Cambridge Bioscience 66305) and developed by exposing to X-ray film (Kodak) after using horseradish peroxidase-conjugated secondary antibodies (Santa Cruz).

### Immunofluorescence (IF)

Cells grown on glass slides or coverslips were fixed with PTEMF (0.2% Triton X-100, 0.02 M PIPES (pH 6.8), 0.01 M EGTA, 1 mM MgCl_2_, 4% formaldehyde). After blocking with 3% BSA, cells were incubated with primary antibodies according to suppliers’ instructions: CREST (Antibodies Incorporated, 15-234-0001), γH2aX (Millipore, 05-636). Secondary antibodies used were goat anti-mouse AlexaFluor 488 (A11017, Invitrogen), goat anti-rabbit AF594, AF488 (A11012, A11008, Invitrogen), and goat anti-human AF647 (109-606-088-JIR, Stratech or A21445, Invitrogen). DNA was stained with DAPI (Roche) and coverslips mounted in Vectashield (Vector H-1000, Vector Laboratories). EdU incorporation and staining was achieved using the Click-It kit (Life Technologies), following manufacturer’s instructions.

### Fluorescence *In Situ* Hybridisation (FISH)

Cells were grown on glass slides, fixed in methanol/acetic acid, then put through an ethanol dehydration series. Pan-centromeric probe (Cambio) was denatured at 85°C for 10 minutes then applied to slides, which were then incubated in a humidified chamber overnight at 37C. The following day, slides were put through a series of washes (one 5 min wash at 37°C in 2xSSC, two 5 min washes at 37°C in 50% formamide/2xSSC, two 5 min washes at RT in 2xSSC). For chromosome painting, paint (Cytocell) was applied to the slide at 72°C for 2 min, then left overnight at 37°C in humidified chamber. The following day, slides were washed once with 0.4xSSC at 72°C for 2 min, then 2xSSC/0.05% Tween at RT for 30 s. After either staining methods, slides were then stained with DAPI, then coverslips with Vectashield were applied and sealed.

### Microscopy

Images were acquired using an Olympus DeltaVision RT microscope (Applied Precision, LLC) equipped with a Coolsnap HQ camera. Three-dimensional image stacks were acquired in 0.2 μm steps, using Olympus 100× (1.4 numerical aperture), 60× or 40× UPlanSApo oil immersion objectives. Deconvolution of image stacks and quantitative measurements was performed with SoftWorx Explorer (Applied Precision, LLC). Analysis was performed using Softworx Explorer.

### Single cell Sequencing

Samples from control and experimentally-induced aneuploid cells were sorted by FACS prior to next-generation sequencing library preparation and data analysis using AneuFinder as previously reported^24,56^ except that the Strand Seq library preparation protocol^57^ was used to create higher complexity libraries. Single nuclei were isolated and stained with 10 μg/mL propidium iodide and 10 μg/mL Hoechst. Single nuclei with low Hoechst/PI fluorescence (G1 population) were sorted into 96-well plates containing freezing buffer using a FACSJazz (BD Biosciences). Pre-amplification-free single-cell whole genome sequencing libraries were prepared using a Bravo Automated Liquid Handling Platform (Agilent Technologies, Santa Clara, CA, USA), followed by size-selection and extraction from a 2% E-gel EX (Invitrogen). Single-end 84 nt sequence reads were generated using the NextSeq 500 system (Illumina, San Diego, CA, USA) at 192 single-cell DNA libraries per flow cell. Demultiplexing based on library-specific barcodes and conversion to fastq format was done using bcl2fastq (v1.8.4, Illumina). Duplicate reads were called using BamUtil (v1.0.3). Demultiplexed reads were aligned to the GRCh38 reference genome using bowtie (v2.2.4) and only uniquely mapped reads (MAPQ>10) were used for further analysis. Copy number annotation was performed using AneuFinder (v1.4.0). Sequence reads are determined as non-overlapping bins with an average length of 500 or 40 kb, a GC correction is applied, and binned sequences are analysed using Hidden Markov model, or Edivisive to determine the most likely copy number states. To negate the inherent sample variation introduced by sequencing single cells, a stringent quality control step was included that uses multivariate clustering to exclude libraries of insufficient quality. Chromosome copy number is plotted as a genome-wide state with clustering of cells based on the similarity of copy number profiles. To refine the breakpoints of 500 kb bin detected CNAs based on the position of 40 Kb bin analysis we used ClonalMasker to identify the same CNAs between 500 and 40 kb analyses and return the 40 kb breakpoints (see Figure S3e for CNA pileups before and after refinement).

### Removal of clonal and subclonal CNAs

CNAs were considered to be the clonal if they shared the same ploidy and their position in the genome overlapped by at least 50%. Using this criteria, an SCNA was considered clonal if it appeared in more than two of the control (DMSO or siControl) cells. These CNAs were then filtered out from the experimentally-induced CNAs for subsequent analysis. In addition, CNAs that occurred at the termini of chromosomes that were less than 1 Mb, or that occurred within, or in close (<1 Mb) proximity to centromeres were removed to prevent potential repetitive sequence-based mapping artefacts. The boundaries of low resolution (500 kb) CNAs were refined using high resolution (40 kb) CNAs by taking each low resolution CNA and resetting its left breakpoint to be the left breakpoint of the leftmost overlapping high resolution CNA, and similar for the right breakpoint. Both refinement and clonal CNA filtering scripts are available at https://github.com/MBoemo/clonalMasker.

### Measurement of distance to large or giant genes

For small CNAs – the distance from each breakpoint was calculated to the nearest end of the nearest large or giant gene. The closest distance was then plotted. For large terminal CNAs the interstitial breakpoint only was used to calculate smallest distance to the closest large or giant gene. A list of large and giant genes, their lengths and positions was assembled from UCSC genome browser (GRCh38).

### Generation of randomly placed control regions

Random genomic coordinates and 1 Mb intervals were generated using the bedtools random script from the BEDTools software^58^. 500 random coordinates were generated, 470 and 376 of which could be mapped to the genome during analyses of gene expression and replication timing respectively.

### RNA isolation, sequencing and analyses

Total RNA was extracted (RNeasy kit, Qiagen) from BJ and RPE1 cells, treated with either DMSO or aphidicolin, from three biological replicates. RNA quality was analysed on Tapestations, with RIN numbers were above 7.0. Library preparation was performed using the Lexogen 3’ tagSeq kit. RNA sequencing was performed by Barts and the London Genome Centre on the Illumina NextSeq 500 platform, generating on average ~1 million single-end reads of 76 bp in length per sample. Raw reads were mapped to the human genome (hg38, Genome Reference Consortium GRCh38) using HISAT2^59^. Number of uniquely aligned reads (q > 10) to the exonic region of each gene were counted using HTSeq^60^ based on the GenCode annotation release 29. Only genes that achieved at least one read count per million reads (cpm) in at least three samples were kept and a log2 transformed cpm expression matrix was subsequently generated. Differential expression analysis was performed using the ‘limma’ R package^61^. In order to analyse total gene expression levels at aCNAs we generated a 2 Mb window of analysis centred around the putative breakpoint. The average expression of all genes in that window was summed. These values were then plotted in categories of small (<20 Mb) or large (>20 Mb) gain and loss CNAs. For large and giant gene expression analysis we used the mean gene expression from three biological replicates.

### Replication Timing

For replication timing both single and bulk G1 cells serving as controls were sorted. For experimental samples we sorted various S-phase populations based on DNA-content (late G1/early S; early S/mid S; mid S/late S; late S/early G2). Single-cell sequencing was performed and reads were processed as described above. The replication timing method as described by Takahashi^34^ was then applied. Uniquely mappable sequence reads were binned into 1 Mb bins. Read counts per bin were then determined and converted into a read proportion across all reads. Bins with a read proportion lower than the 0.1 quantile were excluded. Quantiles were calculated separately for the autosomes and X-chromosome in male samples (the Y-chromosome was not included, neither were aneuploid chromosomes). To correct for variable mappability median centering was applied using the data from G1 cells as a reference. Correction factors were calculated by dividing the average read proportion by the proportion of each bin. Bin proportions were then multiplied by the correction factors to yield a corrected average proportion per bin. To determine whether a bin was replicated or not a quantile cut-off was applied per S-phase population to reflect the fraction of the genome expected to be replicated at that moment (i.e. 0.125, 0.375, 0.635, 0.875 for the population mentioned above). Bins with read proportion greater than the quantile cut-off set per S-phase population were designated as ‘replicated’ (blue in the plots). Conversely bins with proportions below the quantile cut-off were designated ‘unreplicated’ (yellow in the plots).

### Replication timing Factor

For each bin, we determined whether the genome was replicated or not (1= replicated, 0= not replicated), in each of the single cells, at each of the four replication phases (early S phase, mid S phase, mid-late S phase and late S phase). We then summed the values across all replication phases, creating the replication factor value, where higher values represent earlier replication. We then categorised the range of replication factor values in four quartiles. Lowest quartile represents late replicated bins and highest quartile indicates bins that were replicated in early S phase, with mid-early and mid-late quartiles in between the two. To calculate large gene replication timing, we took the average replication factor for the two genomic bins that covered that large gene. For analysis of RT of CNA breakpoints, for small CNAs – the distance from each breakpoint was calculated to the nearest end of the nearest large or giant gene. The closest distance was then plotted. For large terminal CNAs the interstitial breakpoint only was used to calculate smallest distance to the closest large or giant gene.

### SNP 6.0 analysis

SNP 6.0 analysis was performed by Aros AB (Denmark) using a Genome-Wide Human SNP Array 6.0 (Affymetrix) on RPE1 cells and two edited RPE1 lines derived from the parental RPE1 line. Data were analysed in the Chromosome Analysis Suite (CAS, Affymetrix). Data were transformed from global references obtained from signals in the CAS normalised reference library.

### Statistical analysis

Unpaired *t*-test or one-way ANOVA Kruskal-Wallis test with post-hoc Dunn’s correction were used to test for levels of significance using either Excel or Prism (GraphPad). Asterisks have been used to denote the significance value between experimental conditions adhering to the following nomenclature: p<0.05 (*); p<0.005 (**); p<0.0005 (***); p<0.00005 (****).

---

**Figure S1:**
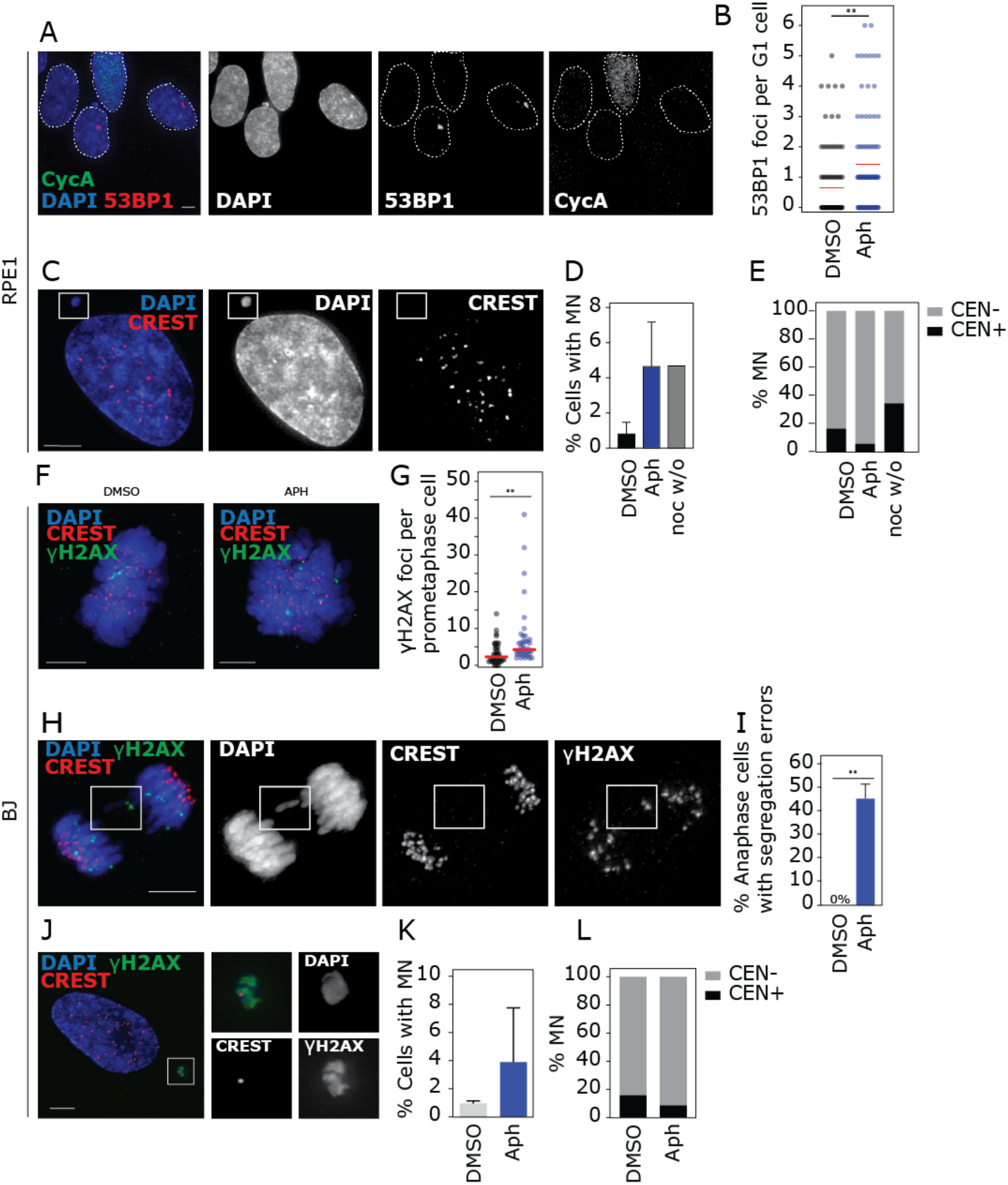
(relating to Figure 1): (**A**) Immunofluorescence image of RPE1 interphase cells, stained for 53BP1 bodies in G1 cells; Cyclin A staining used to identify S and G2 cells. (**B**) Quantification of 53BP1 bodies in G1 cells (n=106 and 107 cells). Red bar indicates mean. (**C**) Representative image of RPE1 cell with a micronucleus. CREST antibody was used to stain for centromeric proteins. (**D**) Quantification of MN rates in RPE1 cells after indicated treatments (three experiments for 24 h DMSO and 24 h aphidicolin, one for 4 h nocodazole and 16 h release; n=1029, 820 or 299 cells respectively). (**E**) Centromere status of MN in RPE1 cells after indicated treatments (n=94, 114 or 38 micronuclei, respectively), as determined by CREST staining. (**F**) Immunofluorescence images of BJ prometaphase cells. (**G**) Quantification of γH2Ax DNA damage foci in BJ cells after indicated treatments (n=66 and 73 cells). (**H**) Representative images of BJ anaphase cells with segregation errors. (**I**) Quantification of segregation errors in BJ anaphase cells. (**J**) Image of BJ interphase cell with micronucleus (**K**) MN rates in BJ cells after indicated treatments (n=743 and 600 cells). (**L**) Centromere status of BJ micronuclei. Statistical tests are unpaired *t*-tests.

**Figure S2:**
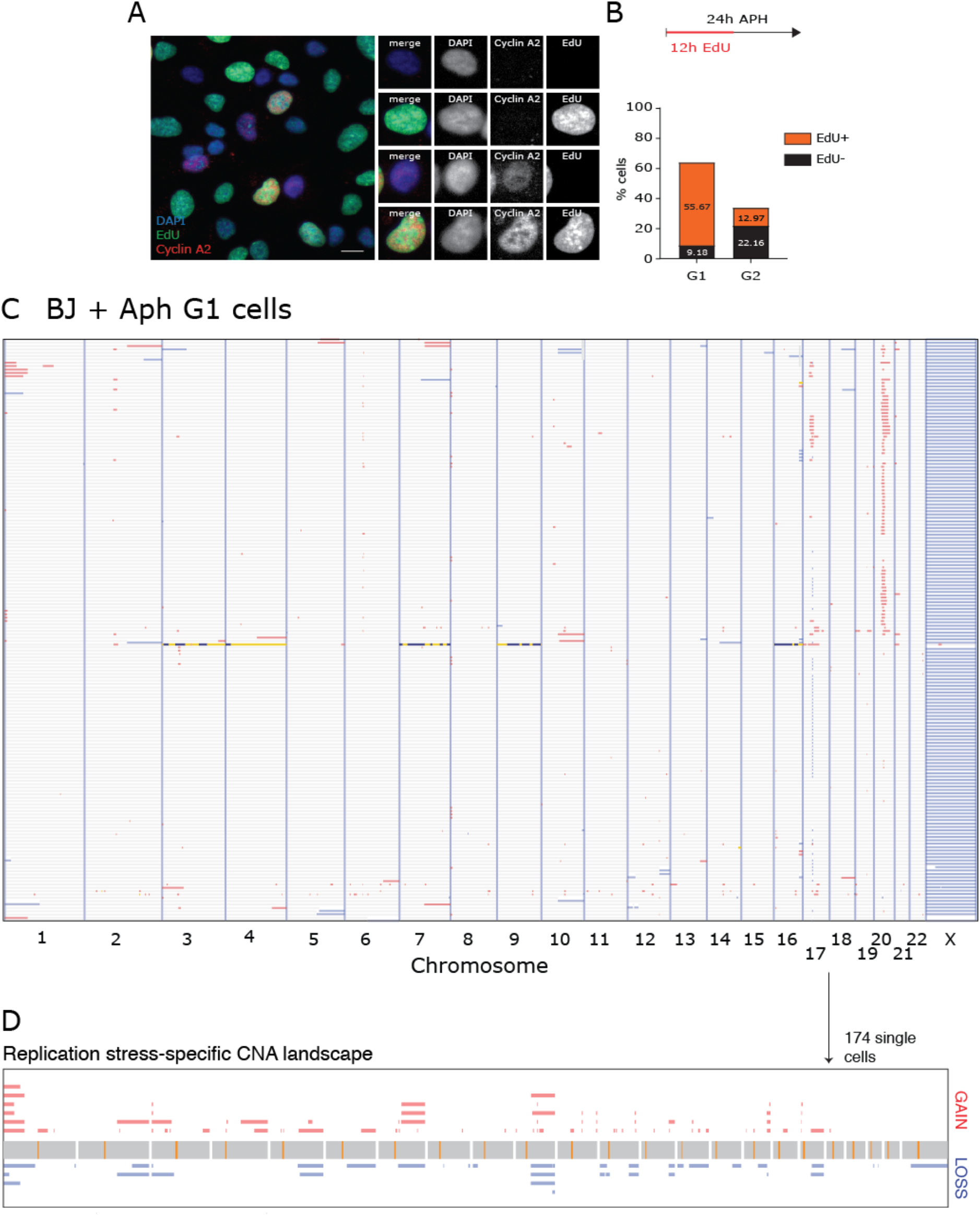
(relating to Figure 1) (**A**) Images of RPE1 cells stained for Cyclin A and EdU incorporation.(**B**) RPE1 cells were treated with DMSO or aphidicolin for 24 h, including a 12 h pulse of EdU as indicated. EdU incorporation into Cyclin A positive (S/G2) or negative cells (G1) was then quantitated from immunofluorescence microscopy images (n=185 and 235 cells). (**C**) Heatmap of single cell sequencing data from 174 BJ cells after 24 h treatment with aphidicolin. (**D**) Diagram summarising all RPE1 aCNAs taken from (C) after removing clonal events (see **Methods**).

**Figure S3.**
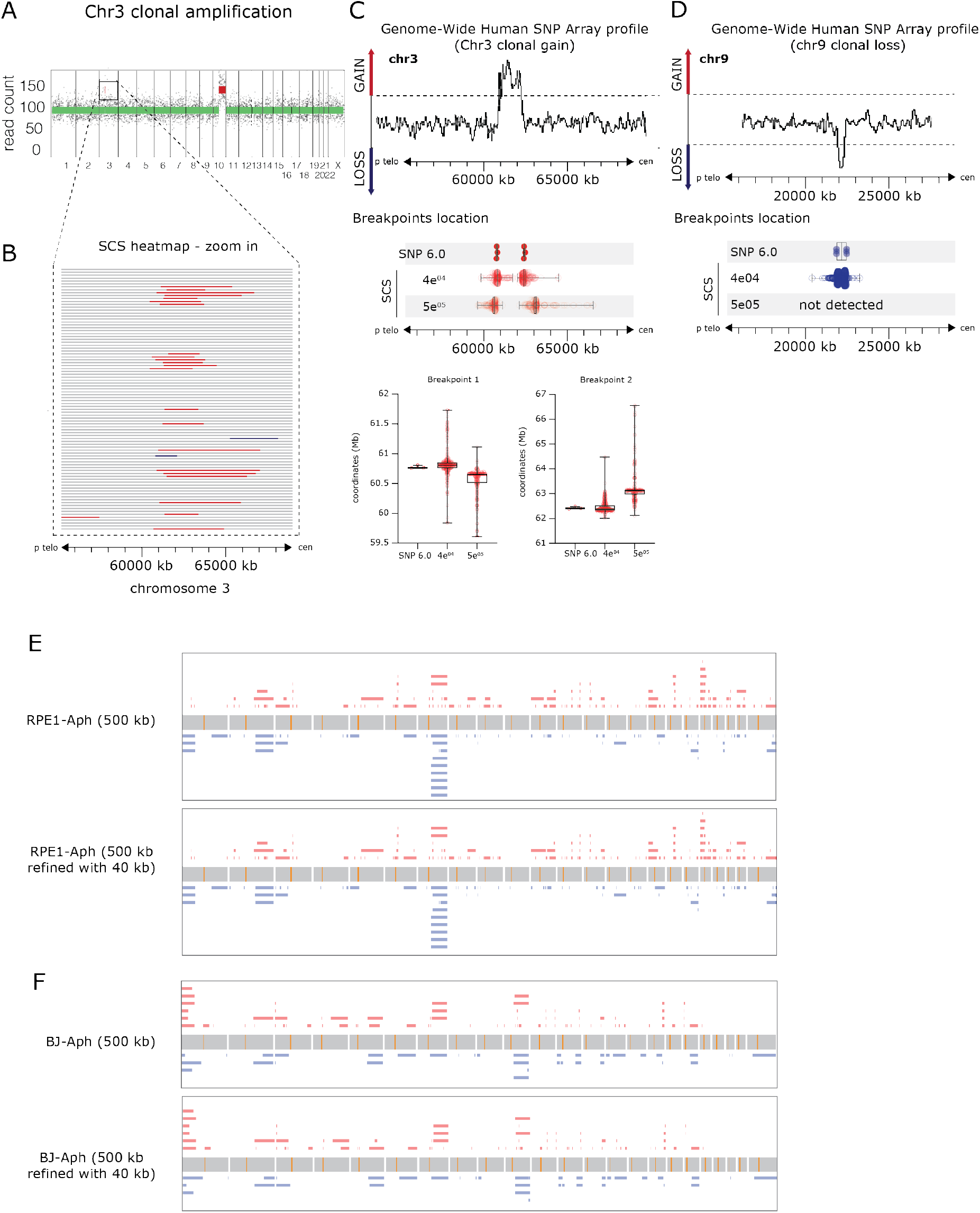
(related to Figure 1): (**A-C**) Schematic indicating SNP 6.0 array verification of clonal focal amplification on chromosome 3 in RPE1 cells, and comparison between 500 kb and 40 kb binning CNA detection accuracy. The three SNP 6.0 datapoints indicate CNA positions determined from three independent RPE1 cell line clones analysed in parallel. (**D**) Detection of ~ 0.5 Mb clonal loss with 40 kb but not 500 Kb binning CNA detection. (**E**) Pileups of all detected aCNAs (after removing clonal events, centromeric and telomeric events (see **Methods**)) before (top panels) and after (lower panels) refining breakpoint position using 40 kb calls (see **Methods**).

**Figure S4.**
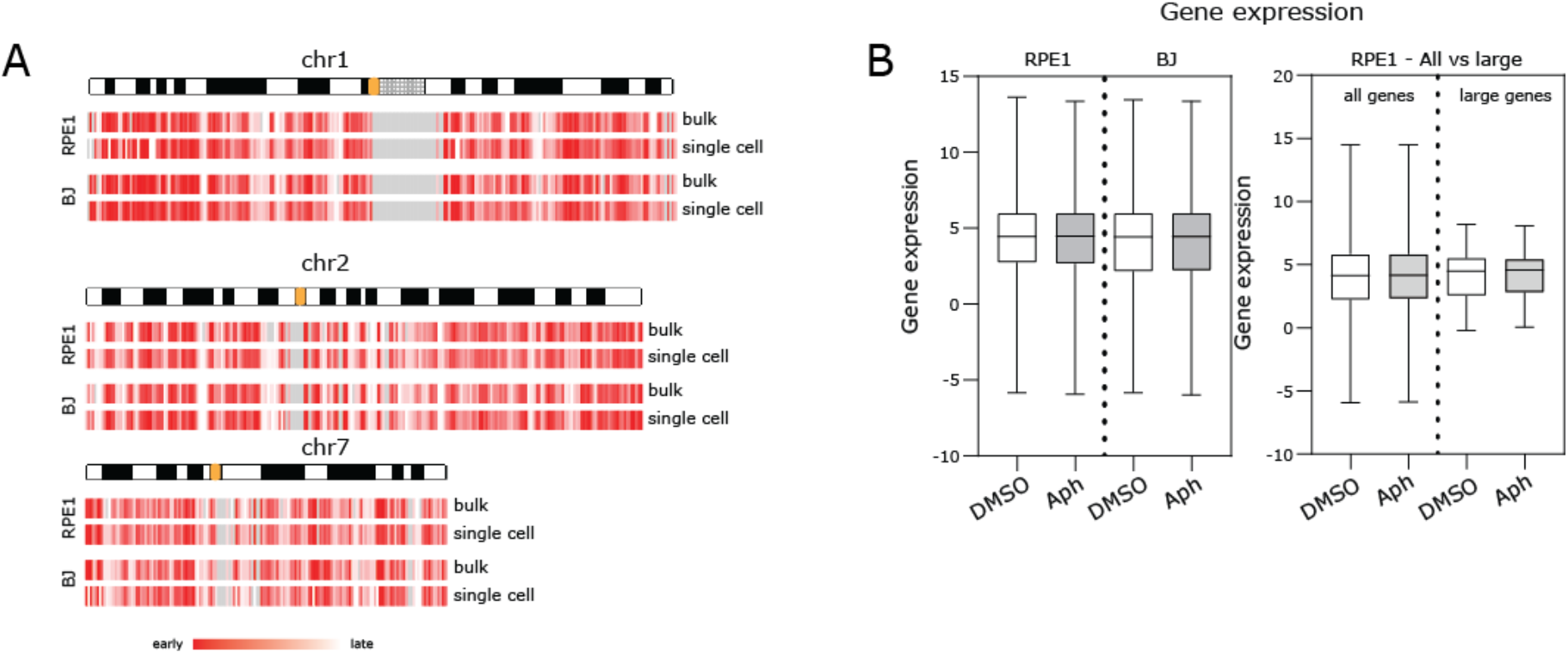
(related to Figure 2): (**A**) Comparison between single cell and bulk replication timing for RPE1 and BJ cells for three selected chromosomes. (**B**) Left side: Total gene expression in DMSO vs aphidicolin for RPE1 and BJ cells; right side: expression in RPE1 cells treated with DMSO vs aphidicolin for all genes vs only large genes (>600 kb).

**Figure S5:**
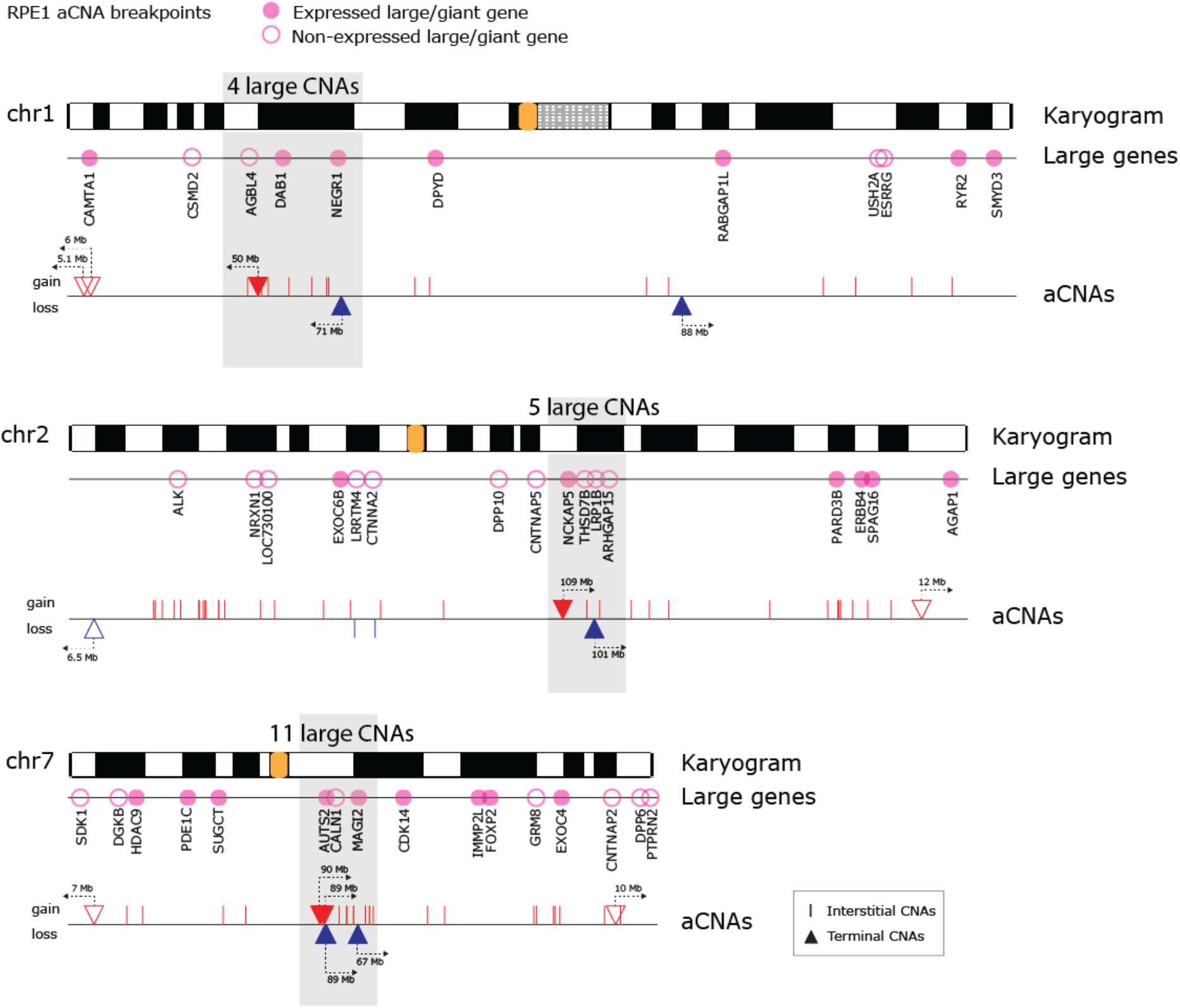
(relating to Figure 3): Karyograms of chromosomes that undergo recurrent large terminal aCNAs (chr 1, 2 and 7) with aCNAs and large/giant genes indicated. Expression of large/giant genes from RPE1 cells is indicated by filled pink circles.

**Figure S6:**
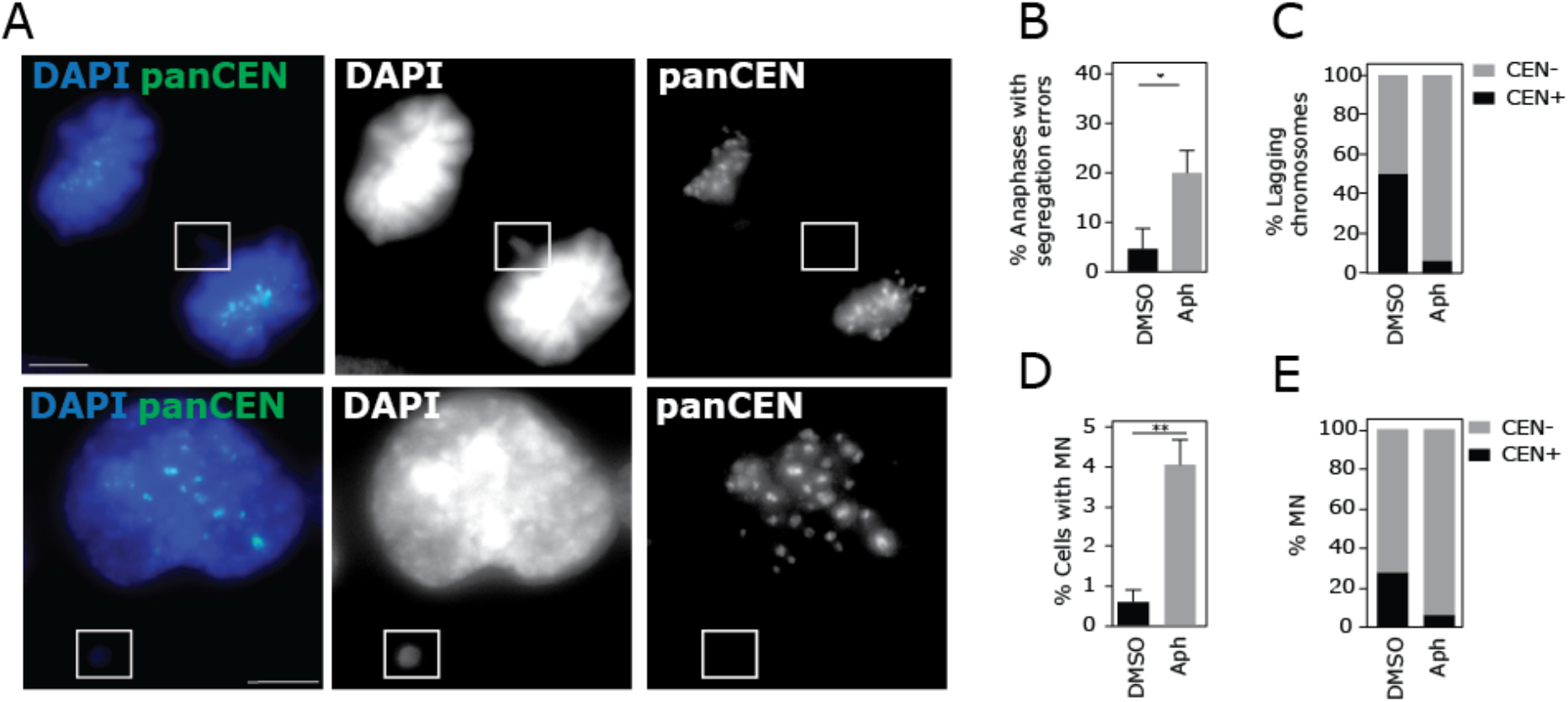
(relating to Figure 4): (**A**) Examples of RPE1 anaphase cells with lagging chromosomes, or interphase cells with MN, stained with pan-centromeric FISH probes. (**B**) Segregation error rates in RPE1 cells stained with FISH probes (n=106-109 cells). (**C**) Centromere status of lagging chromosomes in RPE1 cells as determined by pan-centromeric FISH staining (n=7 or 29 lagging chromosomes). (**D**) MN rates in RPE1 cells (n=725 or 674 cells). (**E**) Centromeric status of MN scored in RPE1 cells after pan-centromeric FISH (n=104-119 MN scored). Statistical tests are unpaired *t*-tests.

**Figure S7.**
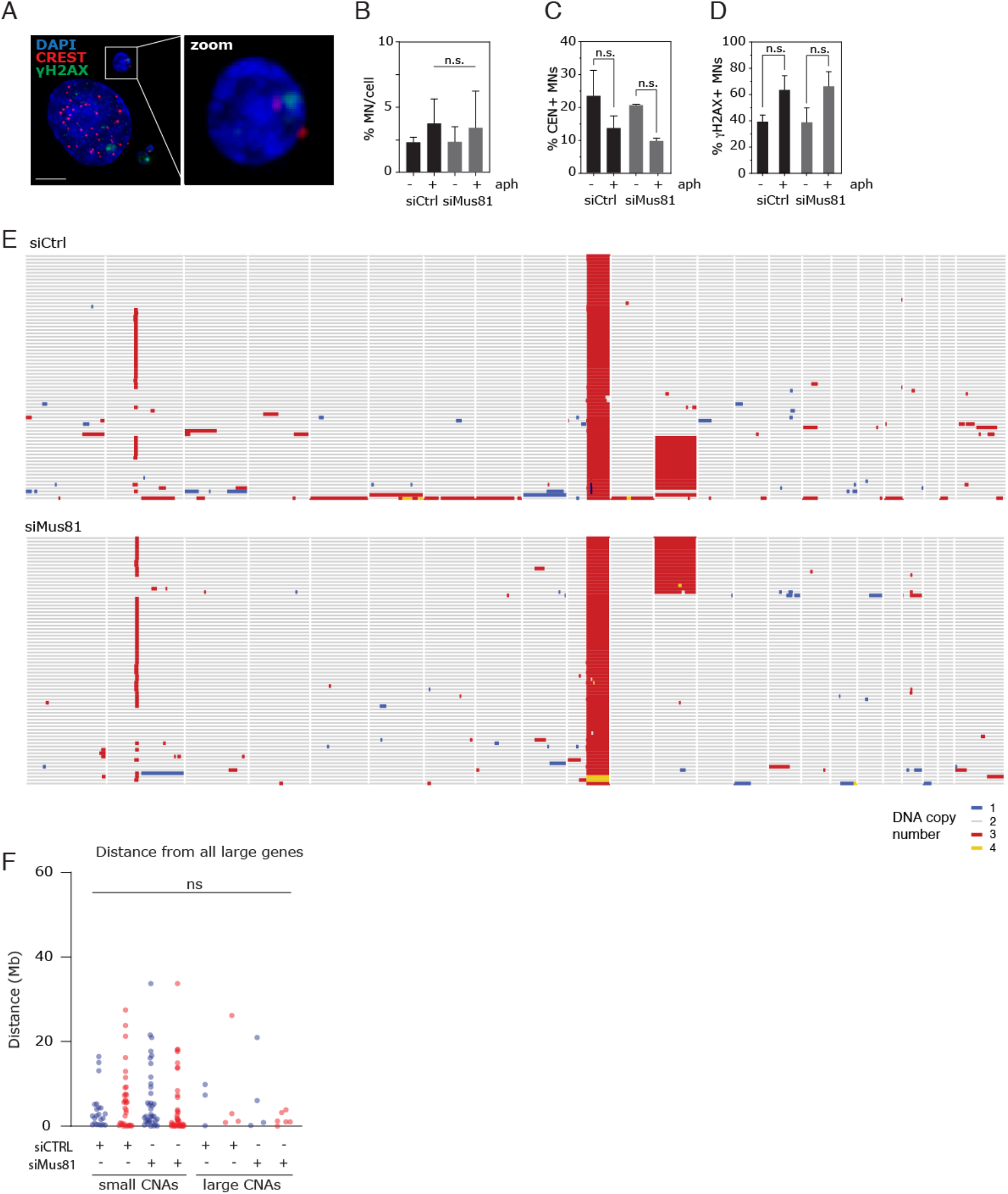
(related to Figure 5): (**A**) Immunofluorescence images of cells with micronuclei (MN). 25 z-stack projections, scale bar is 5 μm. CREST antibodies were used to stain for centromeric proteins and γH2AX antibodies were used to mark DNA damage. (**B**) Quantification of MN per cell (504-792 cells scored in total). (**C**) Quantification of centromeric status of MN. Statistical test was a one-way ANOVA – Dunnett’s test. (**D**) Quantification of γH2AX positive MN(59-113 MN scored in total for C and D). Statistical test was an unpaired t-test. (**E**) Single cell sequencing data from 68 RPE1 cells after 48 h treatment with siControl and 74 cells after 48 h treatment with siMus81. (**F**) Distance from large/giant genes in siControl or siMus81 as indicated. Statistical tests were Kruskal-Wallis tests comparing siMus81 with siControl in each case.

